# Structure and mechanics of the human Nuclear Pore Complex basket

**DOI:** 10.1101/2022.10.20.513091

**Authors:** Anthony Vial, Luca Costa, Patrice Dosset, Pietro Rosso, Gaëlle Boutières, Orestis Faklaris, Heiko Haschke, Pierre-Emmanuel Milhiet, Christine M. Doucet

**Affiliations:** CBS (Centre de Biologie Structurale), Univ Montpellier, CNRS, INSERM, Montpellier, France; MRI, Biocampus, University of Montpellier, CNRS, INSERM, Montpellier, France; Bruker Nano GmbH, Berlin, Germany

## Abstract

Nuclear pore complexes (NPCs) are the only gateways between the nucleus and cytoplasm in eukaryotic cells. They restrict free diffusion to molecules below 5 nm while facilitating the active transport of selected cargoes, sometimes as large as the pore itself. This versatility implies an important pore plasticity. Recently, cryo-EM and AI-based protein modeling revealed with acute precision how most NPC constituents are arranged. But the basket, a fish trap-like structure capping the nucleoplasmic side of the pore, remains the missing piece in this puzzle. Here by Atomic Force Microscopy (AFM) coupled to Single Molecule Localization Microscopy (SMLM) we revealed that the basket is very soft and explores a large conformational landscape: apart from its canonical shape, it dives into the central pore channel or opens, with filaments reaching to the pore sides. Our observations enlighten how this structure can adapt and let morphologically diverse cargoes shuttling through NPCs.

## Introduction

In eukaryotic cells the nuclear envelope (NE) isolates the genome from the rest of the cell. This physical barrier is composed of two biological membranes: the Outer Nuclear Membrane (ONM), that is continuous with the endoplasmic reticulum, and the Inner Nuclear Membrane (INM), which harbors a unique integral protein composition (*1*). Nuclear Pore Complexes (NPCs) are the only gateways between the cytoplasm and nucleoplasm. Their primary function is to ensure the permeability barrier: they restrict passive diffusion to small molecules below 40 kDa, yet allowing regulated transport of larger protein assemblies, including the export of mRNAs (*2*). Very large MDa molecular assemblies can be transported through NPCs in their native state, such as proteasomes or viral capsids for instance (*3–6*). As a matter of fact, NPCs are the largest protein complexes found in eukaryotic cells, with a diameter around 100 nm in humans (*7–10*). They are composed of over 1000 polypeptides for a total size of 110 MDa in humans (*7*). With their 8-fold rotational symmetry, NPCs are composed of about 30 different proteins called nucleoporins (Nups), present in multiples of 8 copies per pore. The NPC scaffold is made of three stacked rings embedded in the NE, and a lumenal ring located in the perinuclear space. The central channel is filled with nucleoporins rich in FG-repeats that form a hydrogel whose physico-chemical properties rule the permeability barrier (*11*). This whole structure is capped by cytoplasmic filaments on one side, and a nuclear basket on the other (Figure 1A).

**Fig. 1.**
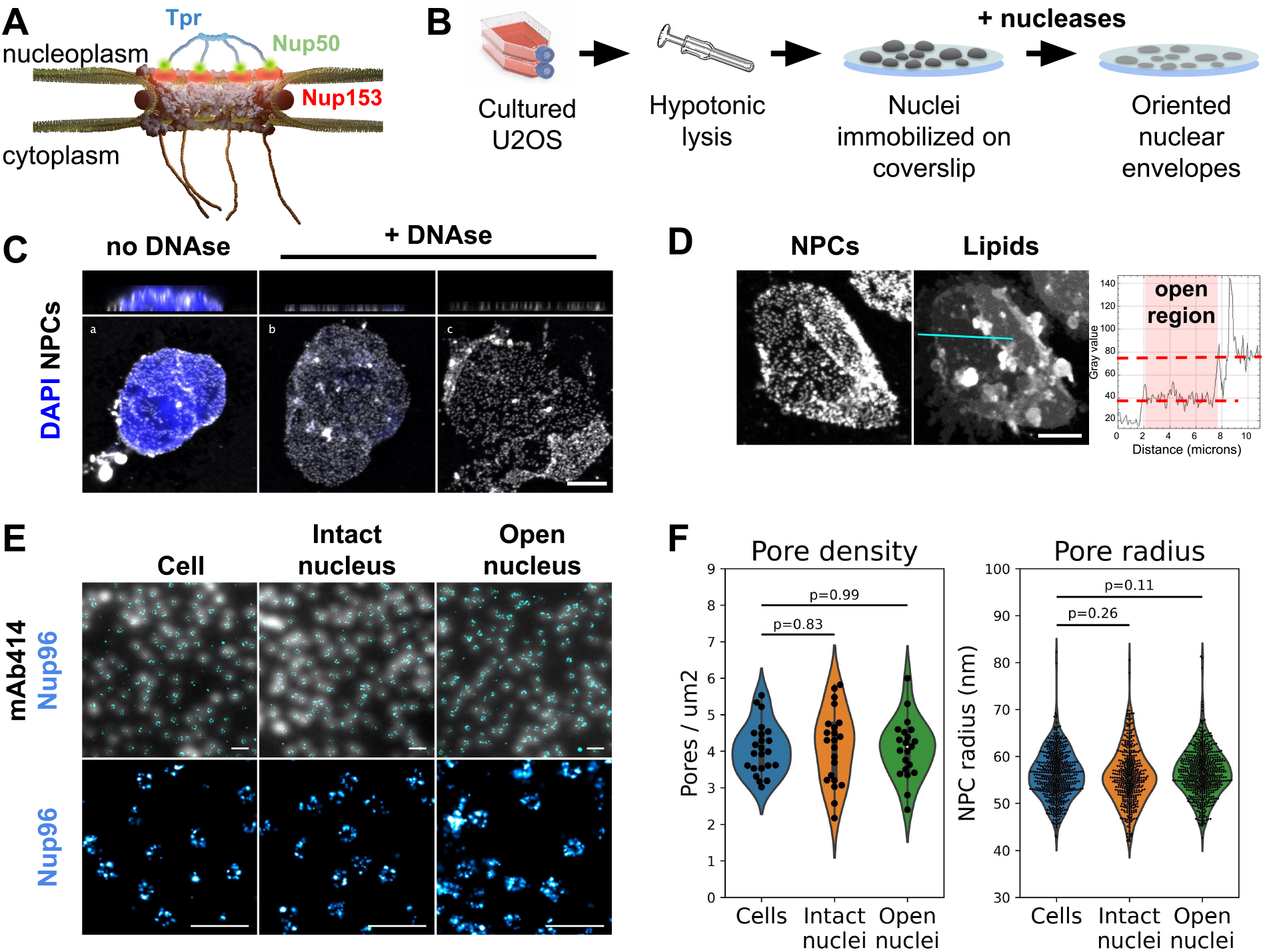
The nuclear membrane is preserved during NE preparation. **(A)** Schematics of the human Nuclear Pore Complex, with emphasis on basket organization. **(B)** Schematic procedure for Nuclear Envelope preparation from cultured U2OS. **(C)** 3D-confocal imaging of nuclei isolated from U2OS and treated or not with nucleases. The chromatin was labeled with Hoechst (blue) and NPCs with WGA (white). Top panels show xz sections; bottom panels show the lower nuclear membrane. Scale bar is 5 μm. **(D)** Confocal imaging of an open nucleus. NPCs are labeled with WGA-AF594, lipids with DiOC6. Scale bar is 5 μm. The right panel shows the intensity profile of the lipid dye along the cyan section. **(E)** Nuclei and nuclear envelopes were extracted from U2OS/Nup96-SNAP. Intact cells, intact nuclei or open nuclei were labelled with mAb414-AF594 and AF647-SNAP ligand. Diffraction-limited (mAb414) and STORM images (Nup96-SNAP) were acquired in TIRF illumination; scale bars are 2 μm (top) and 500 nm (bottom). **(F)** Pore density was measured from confocal images (shown in figure S2C, n>20 nuclei); NPC radii were measured from STORM images as exemplified in E (n≥3 nuclei).

Years of intense structural investigations have gradually revealed the molecular architecture of the human NPC scaffold. Cryo-electron microscopy (cryo-EM) performed on purified NPCs or *in situ* has depicted the arrangement of the luminal, inner, cytoplasmic and nucleoplasmic rings with respect to the NE membrane, at resolutions down to 2 nm (*12–16*). Integrative structural studies have positioned X-ray crystallographic and model structures of individual Nup fragments within the electron-dense volumes. We now visualize with subnanometer resolution how Nups are arranged within the scaffold (*8,12,14–18*). In striking contrast, the cytoplasmic filaments and nuclear basket are systematically absent from cryo-EM maps. One interpretation is that these domains are highly flexible, leading to a continuum of conformations that cannot be resolved by cryo-EM. While a near atomic composite model of NPC cytoplasmic face was recently published (*12*), the nuclear basket is notoriously the last missing piece in the puzzle. From scanning electron micrographs taken in the 90s (*19*) we know that the basket is made of eight filaments attached at their base to the NPC scaffold and joining together at their distal ends. The resulting structure appeared as a fish trap-like net, hence its name - the basket. In higher eukaryotes, this structure is mainly composed of three Nups: Nup153 and Nup50, located at the base of the basket, and Tpr which contains four predicted coiled-coil domains and is the major constituent of the filaments (*20*). During their nucleo-cytoplasmic shuttling, cargoes have to pass through the basket, which caps the nucleoplasmic opening of the channel. It is hard to envision how very large cargoes, such as intact viral capsids (*5*), can make their way in this intricate structure. Indeed, such large cargoes almost fill the entire space within the central pore channel, and their release in the nucleoplasm must involve some opening or disassembly of the basket. As a matter of fact, while most components of the scaffold are very stably associated, several Nups nevertheless can exchange from the NPC (*21*). It is in particular the case for Nups that connect core NPC components together (*22*) and their high exchange rate was proposed to favor structural adaptation of the NPC during cargo translocation for instance. Likewise, components of the basket are highly dynamic: they exchange rapidly with the NPC, indicating a continuous turnover between NPC-bound and free Nups (*21–24*). Moreover, serial analysis of thousands of NPCs imaged by Single Molecule Localization Microscopy (SMLM) show that the main basket constituent, Tpr, exhibits a blurry spatial distribution while other components show a better-defined localization (*25*). However, most studies cited above rely on ensemble or averaged structural data. It is thus not clear how structural dynamics and/or variability of the basket translate at the single pore level. Therefore, to explore different basket conformations, we used here Atomic Force Microscopy (AFM) that does not rely on averaging.

Thanks to its very high signal to noise ratio, AFM provides sharp topography images of single particles and operates in liquid, making it an asset to study heterogeneous samples in physiological conditions (*26*). In AFM, samples are probed by a nanometric tip located at the end of a cantilever. Tip-sample interactions induce deflection of the cantilever, which can be detected and analyzed to reconstruct the topography of the sample surface and can also provide mechanical information. Furthermore, combined with super resolution fluorescence microscopy, molecular mapping can be correlated with AFM nanoscale topography of biological membranes with lateral resolutions up to 10-30 nm (see a review in (*27*) and a study carried out in our group (*28*)). AFM thus appears as a method of choice to observe and describe at high resolution the conformational states of NPCs. However, this technique requires direct accessibility to the probed surface. In the context of NPC study, it means to isolate nuclei and purify NEs. This technique has been developed on *X.laevis* stage-VI oocytes, since their nuclei are large enough (~300 μm in diameter) to be manually teared open (*29*). High resolution images of the cytoplasmic and nucleoplasmic sides of *Xenopus* NPCs have revealed a certain degree of architectural plasticity - their morphology changes in response to different physiological stimuli or addition of transport receptors (*24*, *30–37*). However, these works did not provide an extensive description of the conformational landscape of NPC baskets. In other words, it is not clear whether baskets can adopt a finite number of conformations or if it is a flexible structure that can freely explore a continuum of conformations. Considering the crucial role of NPCs in cellular homeostasis and the fact that they are associated with several diseases (*38–41*), there is a real interest in elucidating the structure of human nuclear pores and understanding their plasticity. Indeed, even though NPCs are highly conserved among eukaryotes, there are compositional and/or structural differences between species (*13*, *42*). Moreover, oocytes are in a particular metabolic state, and this translates into structural and functional specificities even in NPC organization: NPCs are ~10 times denser than in somatic cells, and their structure and size evolves during gametogenesis (*43*). A few studies have been performed on biochemically extracted human NEs (*44*, *45*) but none of them focused on NPC baskets.

We established a biochemical procedure to open nuclei isolated from cultured U2OS cells and imaged by AFM the nucleoplasmic face of hundreds of NPCs in their physiological environment. Thorough statistical analysis showed that the basket largely explores the pore central channel rather than systematically protruding into the nucleoplasm. We also located Tpr within the basket by correlating AFM with super-resolution microscopy and found pores with open baskets, *i.e*. with Tpr-containing filaments reaching towards the membrane around the pore. Comparing the topography of normal NPCs and NPCs devoid of basket demonstrated that the basket contributes to an important volume in the central channel of NPCs, and to a lesser extent at the surface of the scaffold rim. Finally, we found that the basket filaments are very soft, an undeniable asset for structural plasticity. Our findings explain why cryo-EM cannot currently decipher the basket structure but most importantly, they enlighten how it can adapt to cargoes of different morphologies and provide a flexible frame to support nucleo-cytoplasmic transport.

## Results

### Preparation of human Nuclear Envelopes (NEs) with preserved nuclear membrane organization

It appeared recently that biochemically purified NPCs are narrower than their counterparts imaged *in situ* (*8*, *13*). This is arguably due to a loss of tension in the NE. To avoid this artifact, we adapted a classical biochemical procedure to purify NEs (*46*), applying it to nuclei immobilized on glass in order to preserve their spatial organization, orientation and tension. Briefly, a hypotonic cell lysis is carried out using a Dounce homogenizer. The purified nuclei are immobilized on glass coverslips, then treated with a mixture of RNase and DNase in a hypotonic buffer (Figure 1B, see methods for more details). A fraction of nuclei spontaneously opens during this step (Figure 1C). As nuclei are immobilized on glass prior to treatment, when they open, the top NE sheet is removed, leaving the bottom layer intact with its INM facing up. Samples were then fixed and fluorescently labeled for correlative AFM / fluorescence studies. Since nuclei are open, labeling can be performed without detergent, keeping the nuclear membranes intact. We demonstrated that the successive biochemical treatments indeed preserve the nuclear envelope and nuclear pores. While nuclease treatment completely digests chromatin in most nuclei (Figure S1A), it results in different configurations: some nuclei remain apparently intact (Figure 1C-b) while a majority have a fragment of NE ripped off (Figure 1C-c). The areas where one or two layers of NE are present are easily delineated thanks to the two-fold difference in NPC density (figures 1C, S1B) or lipid staining (Figure 1D). Some nuclei expose a well circumscribed central hole; in these cases the membrane organization is well preserved, based on the homogenous lamin-B1 staining (Figure S1C-e,f). These nuclei, easily recognizable, were selected for further studies. On the contrary, some nuclei have large sections of NE ripped off; in those the organization of INM was disrupted, as shown by lamin staining (Figure S1C-c,d), and they were discarded.

Regarding NPC organization, pore densities measured from cultured cells and purified nuclei - treated or not with nucleases - are similar (Figure 1F & S1B). Moreover, direct Stochastic Optical Reconstruction Microscopy (*d*STORM) imaging on NPCs allowed us to measure their diameters (Figure 1E-F). For this, we used a modified U2OS cell-line whose endogenous nucleoporin Nup96 is fused to a SNAP-tag (*47*). Nup96 is a main constituent of NPCs scaffold and its distribution is thus representative of the nuclear pore structure. We labeled intact cells and nuclei - intact or open - with an AlexaFluor 647 SNAP-ligand and performed *d*STORM imaging. The diameter of Nup96-SNAP rings measured in the three situations shows no significant differences (Figure 1F). Conservation of pore density and diameter suggests that adhesion of the NE prior to its rupture preserved its tension, keeping NPCs in their physiological shape. In conclusion, we developed a method to prepare flat human NEs, free of chromatin, with areas of INM accessible to the AFM tip. This procedure preserves membrane integrity and nuclear pore structure.

### Topography of the NPC basket

The notorious absence of the pore basket in averaged structures requires assessing the nucleoplasmic topography of individual pores. Documenting the conformational landscape of the basket can reveal how rapid exchange of Tpr and Nup153 translates in the basket structure. Moreover, this will elucidate to which extent the pore basket can deviate from the canonical fishtrap-like structure into conformations compatible with transport of large cargoes. Figure 2A shows an open nucleus prepared from U2OS cells expressing the transmembrane nucleoporin POM121 fused to GFP (POM121-GFP). Based on pore density, we assessed that this nucleus had a central, well-delineated hole as previously discussed. AFM imaging confirms the presence of this hole surrounded by a bulge of membrane (Figure 2A, right panel). Nuclear pores are clearly delineated by the AFM tip on both the cytoplasmic (black inset) and nucleoplasmic faces (white inset). In order to probe the basket structure, we then focused on the INM. NEs were routinely prepared and labeled with a widely used fluorescent antibody against NPCs recognizing several Nups (mAb414 conjugated to Alexa Fluor 594). The fluorescent pore signal correlates with ring structures in the AFM image (Figure 2B). 250 nm crops around each pore were extracted (examples shown in Figure 2C) and a ring of about 82 nm diameter is visible in all NPCs, reminiscent of previous studies on *Xenopus laevis* NEs (*31–33, 36*) - this is the scaffold. In contrast, the central structure, that corresponds to the nuclear basket, is highly variable: in some cases, filaments are visible (Figure 2C-a,b,d,f,g,j); they are rarely symmetric, but join at their distal ends into a structure reminiscent of the distal ring described in *Xenopus (30–35*). This point either protrudes above the scaffold ring, or stands below the ring level. In other cases, the basket is not clearly depicted and pores appear as an empty barrel. Figure 2E exemplifies these three typical topologies and their radially averaged height profiles (methodological details in the methods section and figure S2A). In the three examples shown in figure 2E, the ring diameter and height are very similar, while the central heights differ a lot. Averaging radial profiles of over 200 individual pores results in a mean profile with the central region lower than the ring (Figures 2D & S2B), far from the canonical protruding basket depicted in textbooks. Consistent with our qualitative description, the standard deviation is higher in the central region of the pore than at the ring level showing more structural variability in the basket than the scaffold. By curiosity we also built an averaged image from all NPC crops (Figure 2F). No basket-like structure is visible in the resulting averaged NPC. This is coherent with our results presented in figure 2D and substantiates that systematic disappearance of basket electron density in high-resolution cryo-EM images is due to high heterogeneity.

**Fig. 2.**
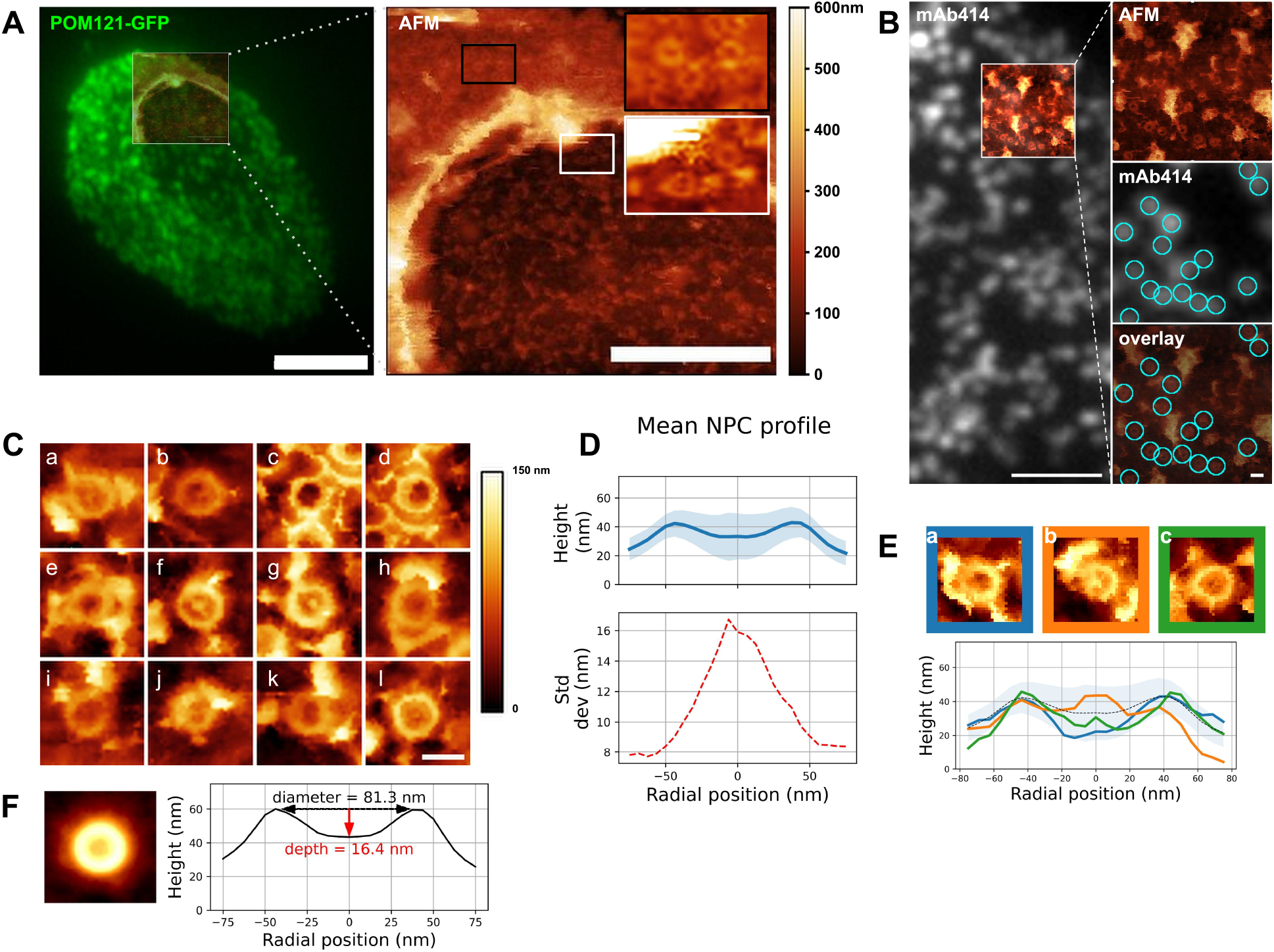
Structure of the NPC basket. **(A)** Correlative fluorescence / AFM image of an open nucleus isolated from U2OS overexpressing POM121-GFP. *left:* TIRF image of the entire nucleus overlaid with the region scanned by AFM. *right:* height image acquired by AFM, encompassing the opening border. Scale bars are 5 μm (left), 2 μm (right). **(B)** Correlative TIRF/AFM image of the inner nuclear envelope of an open nucleus. NPCs are labeled with mAb414 and visualized as fluorescent dots that coincide with ring-like structures in the AFM image. Scale bars are 2 μm and 200 nm. AFM color scale is 0-300 nm. **(C)** Representative samples of human nuclear pores (nucleoplasmic face) imaged by AFM. Scale bar is 100 nm. **(D)** Mean of rotationally averaged NPC height profiles (*n=210*). The standard deviation (shaded area in the top panel) is plotted in the bottom panel. **(E)** Three configurations of NPC nucleoplasmic region and their respective height profile. “Empty” (blue), protruding (orange) and low basket (green). **(F)** (*left*) Average image of 210 NPC crops. AFM color scale as above. (*right*) Height profile of the resulting image.

We then extracted the diameter and depth from each rotationally averaged profile (Figure S2C and Methods). Diameters are normally distributed, with a mean of 81.5 ± 16.3 nm (Figure S2D). Previous AFM studies on *Xenopus* NPCs documented similar pore diameters of around 81-85 nm, for the cytoplasmic side (*24, 31–33, 48*) and 80-88 nm at the nucleoplasmic side when measured at the ridge (*31*, *32*, *48*, *49*). The mean diameter that we measured here from human INM thus falls in the same range. The depth distribution illustrates that the distal point of the basket can sweep across 60 nm in the axial direction (Figure S2E), which is roughly the height of the human NPC scaffold itself (*8*). This suggests that the basket can explore a continuum of conformations. The height distribution is best fitted with two gaussians corresponding to protruding (depth >0) and collapsed (depth <0) baskets. The relative gaussian amplitudes confirm that protruding baskets are a minority.

Altogether our results show that, while the ring structure is rather conserved from pore to pore, the central region corresponding to the basket is extremely variable and a majority of basket structures do not protrude into the nucleoplasm but rather reside in the central pore channel. Considering that Tpr is a main component of the basket filaments (*20*), a quantitative evaluation of Tpr organization within pores, especially when an empty ring is observed, was performed.

### Tpr organization

Tpr is a 267 kDa protein that contains 4 coiled-coil regions in its N-terminal part while its C-terminus is mostly disordered. Coiled-coils are helical domains that can oligomerize and constitute rigid molecular rods (*50*, *51*). In the case of Tpr, the coiled-coils are interleaved with flexible linkers. Tpr thus contains the structural ingredients to provide filamentous yet articulated segments, which may be an asset for the basket to adapt its shape to cargoes.

We labeled Tpr by indirect immuno-fluorescence on purified NEs and performed correlative *d*STORM/AFM. As expected, Tpr signal correlates with NPCs in the TIRF and AFM images (Figures 3A & S3A). This is even more evident in the 3D representation (Figure 3B), where Tpr density precisely coincides with height signal in the AFM images attributed to the basket domain. An ensemble view shows that Tpr localization densities are not homogenous (Figure S3A). Although this information cannot be quantitatively inferred to numbers of molecules, especially with immuno-fluorescence labeling (*47*, *52*), it nevertheless indicates a variability in the number of Tpr filaments bound to NPCs. This is most likely the result of Nup153 and Tpr turnover (*21*–*23*). At the single pore level, several configurations are observed (Figure 3C): in the first example, a canonical basket is visible in the AFM image and a strong *d*STORM signal correlates with the filaments and the distal ring. Importantly in this example, both height and Tpr signals are asymmetric, supporting that Tpr filaments can partially dissociate from the basket, leading to asymmetrical structures. Other NPCs, like the second example, look empty, with no apparent basket structure in the AFM image but Tpr signal is distributed in distinct localizations, located on top of the scaffold ring or even outside of it. Importantly, Tpr signal still colocalizes with filamentous structures visible in the AFM images. This suggests that the basket is in an “open” configuration, with nucleoplasmic filaments attached at their basis to the NPC scaffold, but their distal ends remaining free. Finally, a third example of NPC exhibits some internal density correlated with Tpr but lower than the NPC ring. This configuration looks like a “collapsed” basket, where the filaments may join by their distal ends but are directed inwards the NPC channel instead of protruding towards the nucleoplasm.

**Fig. 3.**
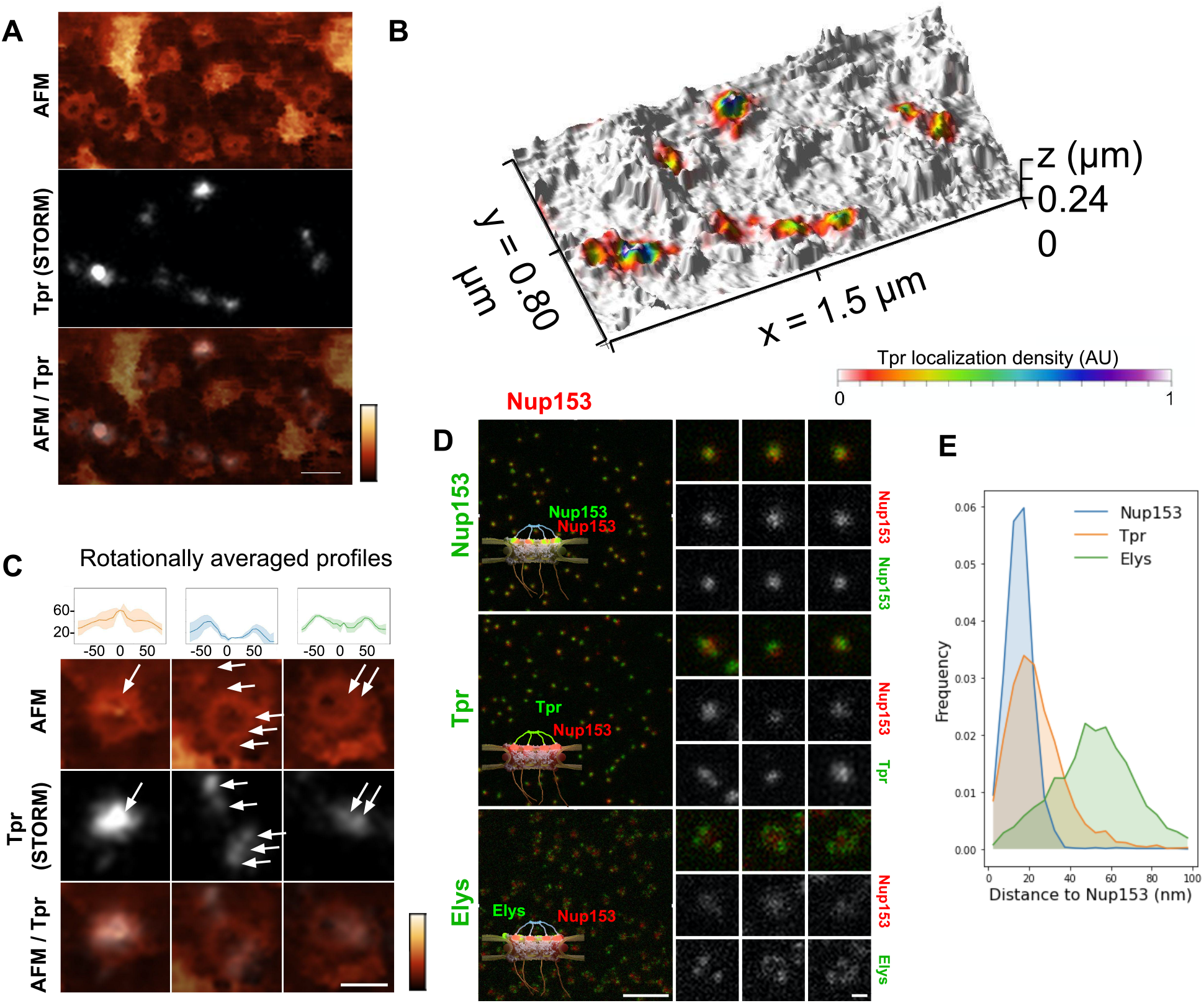
Tpr organization within the NPC basket. **(A)** Correlative AFM/STORM image of an open nucleus prepared from U2OS cells. The sample was immuno-labeled against Tpr. STORM imaging was performed in TIRF illumination. The same area was then imaged by AFM. After reconstruction of the STORM localizations map, the two images were correlated (scale bar is 200 nm). AFM color scale 0-300 nm. **(B)** 3D representation of the correlated AFM/STORM image. **(C)** Three NPCs of typically different topographies are shown in more details, together with their rotationally averaged height profiles (upper panel). Arrows point at Tpr localizations (scale bar is 100 nm). AFM color scale 0-300 nm. **(D)** U2OS cells were fixed and simultaneously immunolabeled against Nup153, Tpr or Elys, with a secondary antibody coupled to STAR-635P (green) and Nup153 coupled to AlexaFluor 594 (red). Samples were imaged by STED microscopy. Individual NPCs are zoomed on the right (scale bars are 1 μm and 100 nm). **(E)** Distance distribution between the AlexaFluor 594 and STAR-635P signals measured from STED images (number of pores >2700).

These images are reminiscent of the three NPC conformations described previously (Figure 2E) and further document the fact that the NPC basket can adopt various configurations. Importantly, as *d*STORM imaging was performed before AFM imaging, the asymmetry observed in Tpr organization in some NPCs is not due to deformation by the AFM tip. However, it could be due to deformations that arose during the biochemical preparation. To check this, we explored the basket structure in intact cells by 2-color STED (STimulated Emission Depletion) microscopy. This confocal-based super-resolution microscopy technique is highly convenient for multi-colour imaging and achieves a typical lateral resolution in fixed biological samples around 40 nm (*53*, *54*). We performed co-immuno-fluorescence in fixed cells against Tpr (coupled to a Star-635P fluorophore) and Nup153 (coupled to an Alexa Fluor 594) to visualize their relative organization. As a control, we used Nup153 co-labeled with Star-635P and Alexa Fluor 594; as expected, Nup153 imaged in both channels renders a very similar signal. Conversely, Elys which is one of the outmost nucleoporins (*25*), is localized as a rim around Nup153 central dot, with some variability in the labeling efficiency (Figure 3D). Regarding Tpr, it usually appears as one - sometimes two - dots that can be well aligned with Nup153 signal or slightly shifted (Figure 3D, middle row). To characterize the relative positioning of these various proteins, we measured distances between Nup153 labeled with Alexa-594 and Nup153, Elys or Tpr labeled with Star-635P (Figure 3E). The distribution shows that a fraction of Tpr is shifted away from Nup153, towards the nucleoplasmic ring where Elys resides. These results obtained on entire cells confirm the variability and asymmetry of the NPC basket observed in AFM images.

Altogether, our AFM and super-resolution microscopy data demonstrate important basket plasticity in the human NPC: Tpr-containing filaments can either protrude towards the nucleoplasm, collapse within the channel or partially disassemble at the level of the distal ring, with filaments pointing outwards from the pore channel axis.

### Contribution of the nuclear basket in the topography of NPCs

The presence of interleaved disordered linkers between its coiled-coil domains presumably allows Tpr to explore a large range of conformations, from a completely extended to a rather globular shape - as predicted by AlphaFold for instance (*55*). It is thus hard to predict how deep the basket could collapse into the pore channel. To understand the contribution of the basket in the overall NPC structure, we imaged pores in open nuclei depleted from Nup153. Indeed, as reported previously (*56*), siRNAs against Nup153 prevent Tpr binding to NPCs. Triple labeling of Nup153, Nup96, and Tpr (Figure 4A) confirms that Nup153 depletion is associated with the absence of Tpr, while Nup96 remains visible (Figure 4A, arrows point at pores without Nup153 or Tpr). It is important to note that siRNAs against Nups only achieve partial depletion (*57*). In the case of Nup153 depletion, it results in a mild decrease in pore density (Figure S4A) and importantly, only a fraction of the remaining pores are completely devoid of Nup153 (Figure S4B), which complicates the interpretation of topological differences observed between depleted and naive nuclei.

**Fig. 4.**
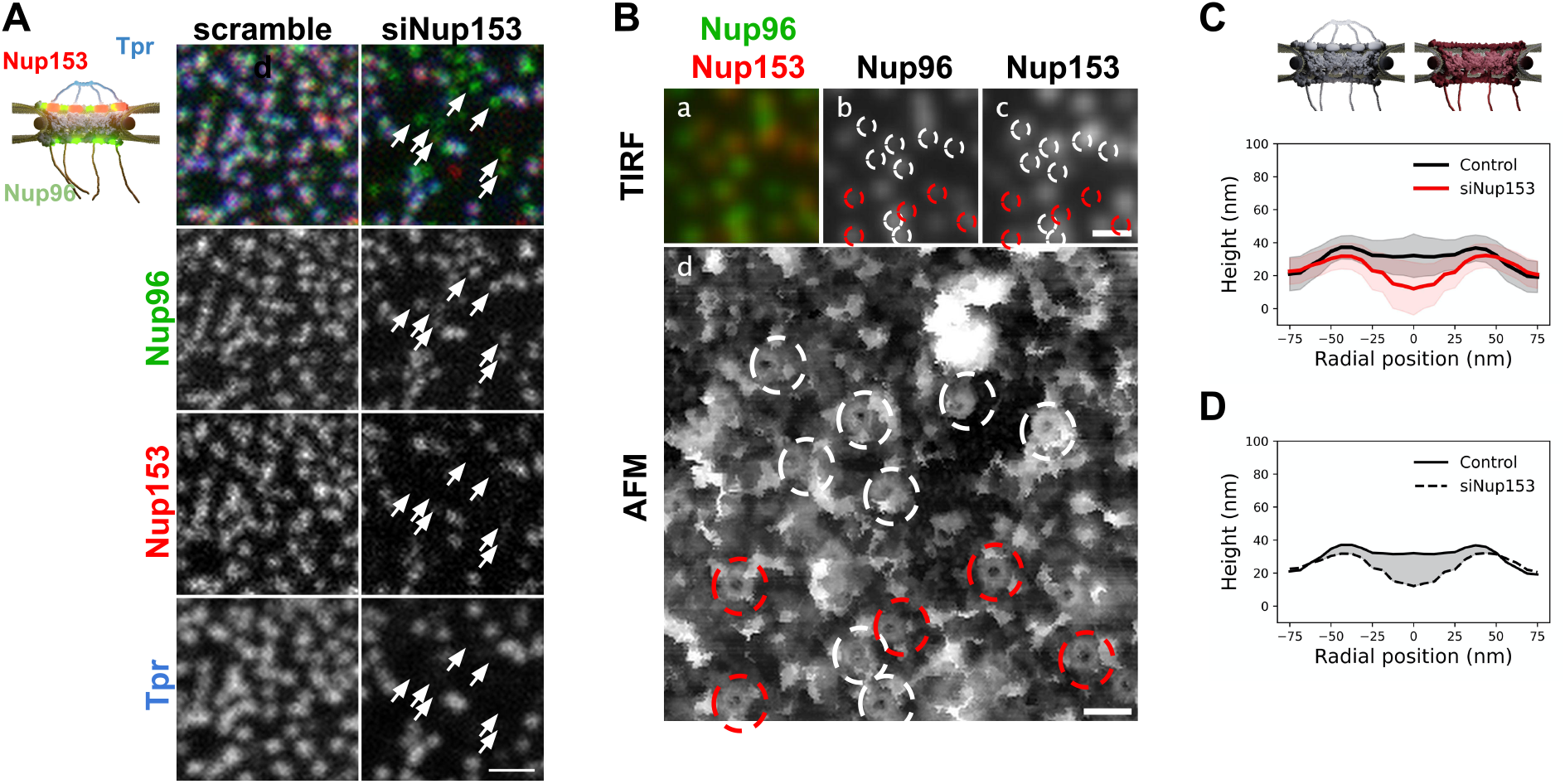
Basket contribution in the topography of human NPCs. **(A)** Stable U2OS cells expressing Nup96-GFP were transfected with siRNAs, scrambled or targeted against Nup153. Cells were then fixed, co-labeled with anti-Nup153 (red) and anti-Tpr (blue) and imaged by confocal microscopy. Arrows indicate pores depleted of Nup153. They are consistently co-depleted of Tpr (scale bar is 1 μm). **(B)** Correlative AFM-fluorescence image of a NE prepared from U2OS/Nup96-GFP cells depleted of Nup153 by siRNA and labeled with anti-Nup153. Panel d shows the ROI scanned by AFM. Top panels show the corresponding Nup96 (b) and Nup153 (c) channels imaged by TIRF. In panels b,c and d, NPCs are circled in white when Nup153 is detected and in red when it is absent or weak. Scale bars are respectively 500 nm (a-c) and 200 nm (d). **(C)** Rotationally averaged height profiles were averaged from over 80 pores imaged from control (black) and siNup153-treated cells (red). Shaded areas are standard deviations. **(D)** The average contribution of the basket in the NPC structure can be envisioned as the volume located between the average surface of control NPCs and of basket-depleted NPCs. A cross-section of this volume is represented as the grey shaded area.

In nuclei with co-labeled Nup96 and Nup153, we can distinguish in correlative AFM / fluorescence images pores that contain Nup153 (circled in white in Figure 4B) or not (circled in red). In the corresponding AFM image, a central topography is more frequent in Nup153-associated NPCs. In contrast, a deep hole is more frequently observed in pores devoid of Nup153. Importantly, even when Nup153 is detected, its levels seem to vary from pore to pore but it is in all cases lower than in mock-depleted cells. To quantify how Nup153 depletion affects NPC topography, we generated average height profiles for pores imaged from Nup153-depleted NEs and compared to naive pores (Figure 4C). Average height measured at the center is dramatically decreased when Nup153 is depleted. The height standard deviations (shaded areas in Figure 4C) partly overlap, reflecting that some pores from depleted cells have similar topographies as compared to control pores. This overlap is likely due to remaining levels of Nup153. One can nevertheless note that very few Nup153-depleted NPCs were seen with protruding baskets (Figure S4C). This conformation may require high levels of Nup153 and Tpr, which is very unlikely to occur in depleted cells. Finally, the very low topographies observed in depleted nuclei probably reflect a total absence of basket. In this case, the AFM tip most likely probes the central FG-repeats. Variability of the central topography may reflect plasticity of the central channel FG-hydrogel. Not to forget also that the central channel is partially occupied by shuttling cargoes, which may contribute to the overall topography and add up to the inherent variability. Anyway, this height difference between control and basket-depleted pores confirms that in most cases, even when the basket shape is not clearly depicted by AFM, it accounts for some molecular density occupying part of the central channel. This average contribution is visualized in Figure 4D, as the shaded area. The height of the scaffold ring is also lower when Nup153 is depleted. This height difference reflects that the basket sits on top of the scaffold. Nup153 is probably present at this location since it interacts with the nucleoplasmic ring (*23*, *58*). Nup153 is mostly intrinsically disordered (*59*) and its structure is likely more fuzzy than the highly ordered components of the nucleoplasmic ring (*8*). This could explain the important volume occupancy at the top of the scaffold.

### Mechanical properties of the basket

Altogether, our data show that the basket can adopt a large range of conformations, such as protruding into the nucleoplasm but also exploring the pore interior. We wondered whether this important structural variability is related to mechanical flexibility. AFM is a very potent technique to measure mechanical properties of soft biological samples, down to stiffnesses of a few pN/nm. Since the imaging mode that we used - Quantitative Imaging (QI, Bruker Nano GmbH, Germany) - is a force curve imaging mode allowing quantitative measurement of mechanical data at each pixel, stiffness maps were generated to understand the mechanical properties of NPCs. Figure 5A shows an AFM height image and its corresponding stiffness map. Rings are seen in the stiffness maps and correspond to NPCs in the height images. Looking at individual pores revealed that stiffness at different regions of the basket varied greatly from pore to pore. To precisely correlate the stiffness to different parts of the NPC, we measured height and stiffness profiles in individual pores along a single line and plotted them on top of each other (Figure 5B). The height profiles (black lines) reveal the scaffold region - the two peaks around ± 42 nm, grey shaded areas - and the basket region - in between these two peaks. In most cases, the scaffold region is the stiffest (in the 4-8 pN/nm range) and the centermost region is the softest (2-5 pN/nm). In a few cases though, the protrusion corresponding to the distal ring coincides with a point of high stiffness (Figure 5B-c). Sometimes the region in between, where the basket filaments must stand, is stiffer (Figure 5B-b&d). To get a general view of NPCs mechanical properties, we generated rotationally averaged stiffness profiles, using the same methodology as for height profiles, and averaged them over 80 individual NPCs (Figure 5C). This shows that on average, the basket region of NPCs is softer than the scaffold. Similarly, stiffness measurements of the NPC nucleoplasmic side in *X. laevis* oocytes obtained by AFM indicated a soft basket, relative to the nucleoplasmic ring (*31*). Of note, our results obtained under mild fixation conditions might overestimate the stiffness of the NPC basket which is expected to rigidify after fixation. In any case, the high standard deviation for stiffness values (red shaded area) underlines the large mechanical variability in the central region of pores. The origin of this variability is not clear but may reside in the transport status of pores: indeed, the density and size of cargoes located in the central channel may influence the stiffness measured in this area.

**Fig. 5.**
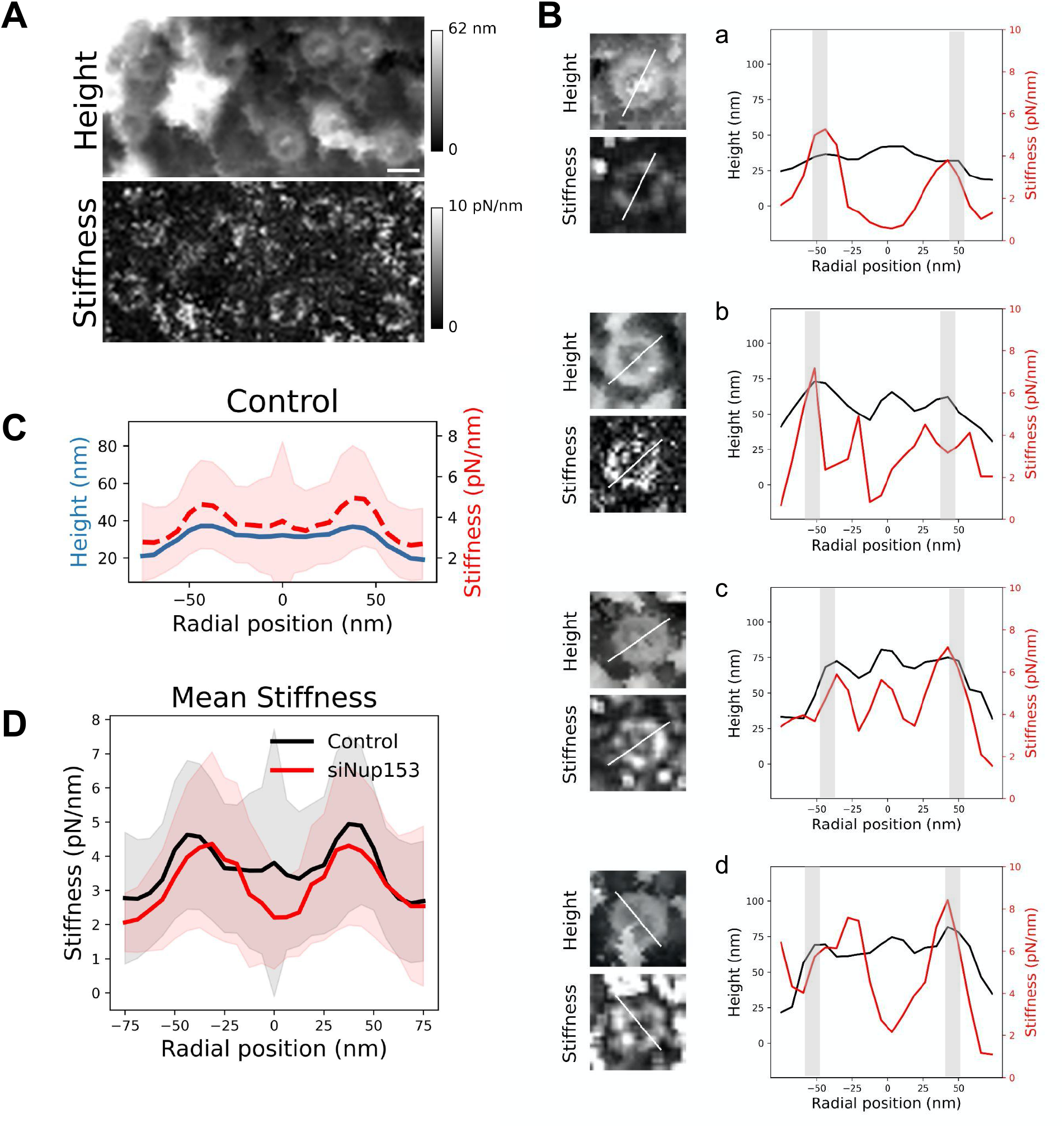
Mechanical properties of the NPC basket. **(A)** AFM Height and stiffness images of NPCs. Scale bar is 100 nm. **(B)** Height and stiffness profiles measured from single pores (along the line depicted on the left panels). The light grey bars indicate the location of the scaffold’s ring. **(C)** Mean of rotationally averaged height (blue) and stiffness (red, dashed) profiles obtained from over 80 individual NPCs. **(D)** Mean of rotationally averaged stiffnesses of NPCs from nuclear envelopes prepared from control (blue profile) or Nup153-depleted (red) U2OS cells. Standard deviations are represented as shaded areas.

Interestingly, when the basket is absent in Nup153-depleted NPCs, the central region of pores is even softer (figure 5D), with a stiffness around 2 pN/nm. In this configuration, the tip is likely directly probing the FG-repeats Nups that fill the central channel and are not shielded by the basket anymore. These proteins assemble as a hydrogel *in vitro (60*) and their stiffness has already been measured by AFM, rendering values in the 5-10 pN/nm range (*60*–*62*), although these may be overestimated in these cases because of the contribution of the underlying substrate. Importantly, we controlled that the basket deformation is not due to the force applied by the tip (Figure S5B).

## Discussion

The huge size combined with their transmembrane nature make NPCs challenging objects to study and in particular at the structural level. Decades of intense integrative structural studies have provided a good understanding of the structure and molecular arrangement of NPC components in the scaffold domain. Yet, the central FG-Nups, cytoplasmic filaments and nuclear basket have been historically missing in these high resolution images, no matter whether they were determined from purified NEs (15, 16, 63) or in situ (8, 13, 64, 65). Recently though, the FG-repeats and cytoplasmic filaments structure and organization have been elucidated. But the basket remains as the last missing piece in the puzzle of NPC molecular organization. Here we show by AFM imaging of human inner nuclear membranes that the basket is highly plastic and can explore an important volume of the central channel of the pore. In particular, the basket filaments can dissociate at their distal ends and open to free the way to large cargoes. When reaching outwards the pore, they may also scan around the pore and facilitate mRNP binding. These findings allow us to propose a model where the conformational flexibility and the dissociation of the distal ends of the basket proteins allow NPC to adapt to cargoes of different sizes (Figure 6). They also give clues as to why high resolution pictures of the basket are systematically missing in cryo-EM maps since it can adopt a continuum of conformations. In addition, the electron density in this region may be much lower than in the scaffold region. Indeed, a rapid calculation based on molecular weights of Nup153, Nup50 and Tpr, and their stoichiometry in human cells (46), indicates that the basket accounts for roughly 13% of the total NPC mass. Yet we show that it can explore a large volume (Figure 4D). Consequently, the electron density from the basket might well fall below the detection limit in cryo-EM.

**Fig. 6.**
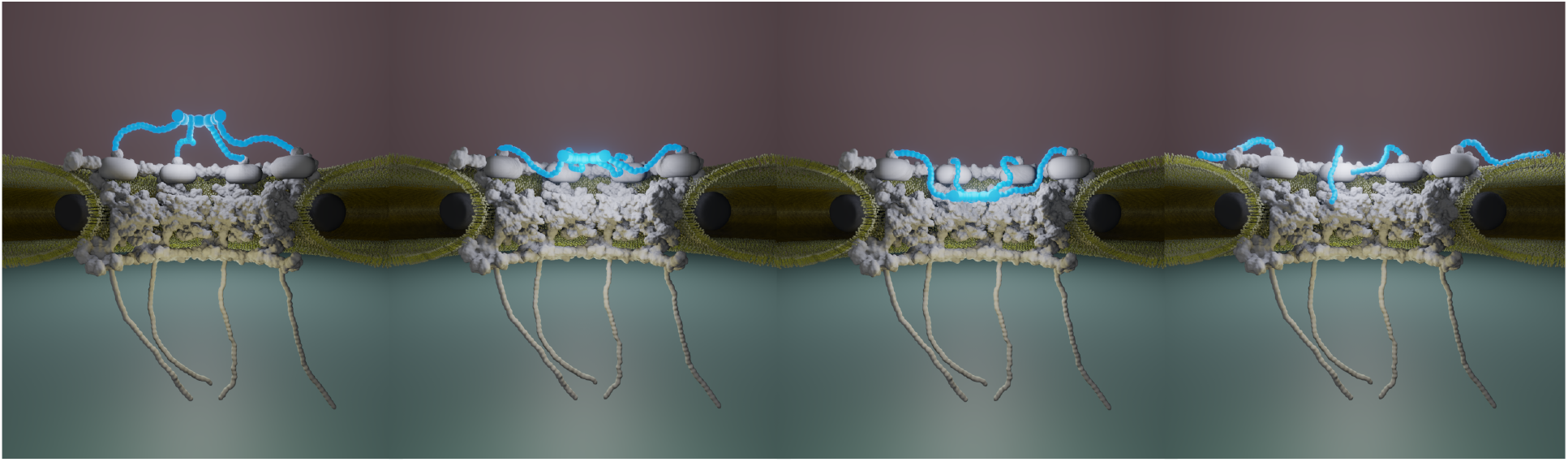
NPC basket variability. This cartoon illustrates a subset of conformations that the basket (in blue) can adopt. The mechanical softness of the basket allows the Tpr-containing filaments to explore a continuum of conformations from an extended configuration, protruding into the nucleoplasm, to a collapsed conformation where the filaments dive into the central pore channel. In addition, the filaments distal ends can dissociate, leading to an open conformation.

These considerations underline the unique possibilities offered by AFM, which does not require averaging and allows to extract mechanical information, to study the structure of the basket. Structural parameters extracted from individual NPCs can then be statistically analyzed. This quantitative approach allowed us to appreciate the plasticity and heterogeneity of the nuclear basket in unperturbed conditions. We thoroughly quantified the topography of hundreds of nuclear pore baskets, and extracted characteristic measurements such as the central depth. Most importantly, we showed that the position of the highest region of the basket can protrude above the scaffold or collapse into the central channel, with positions covering a distance of about 60 nm (Figure S2E). This suggests that the basket can explore a continuous conformational landscape. Being less restricted than other parts of the nuclear pore, the peripheral domains (cytoplasmic filaments and basket) require a lower entropic cost to change conformation. In this regard, they may be an asset for NPCs to adjust their shape.

NPC plasticity had been previously explored by AFM (31, 32, 36, 48, 66–70), mostly on NEs prepared from Xenopus laevis oocytes, for reasons already mentioned: ease of NE preparation and high pore density. Of these, many studies have focused on the cytoplasmic sides of NPCs as the FG-Nups are directly accessible from this side and are not occluded by the nuclear basket. This allowed to study the dynamics and plasticity of the central channel components in response to various signals (calcium, Nuclear Transport Receptors, steroids, apoptotic signals…). It was shown in particular that a mass located at the center of the pore - often referred to as the “central plug” - varies. Because AFM does not offer chemical signatures, the nature and origin of this mass was not clearly attributed. It could be parts of the pore subjected to conformational changes, or cargoes being transported. In the present study, we took advantage of correlative dSTORM / AFM to assign Tpr to specific structures delineated by AFM. This let us unambiguously attribute the basket structure, and was especially helpful to identify unexpected conformations such as collapsed or open baskets (Figure 3C). The ability of the basket to explore a wide volume around its expected protruding position was surprising but this aspect is functionally relevant, as discussed further below. However, as our study only provides a static view, we have to elaborate hypotheses to explain the conformational plasticity of the basket. In a first scenario, filaments would explore nucleoplasmic regions around the pore or fold back inside the channel while remaining attached to the pore scaffold at their basis. This scenario is plausible given the very low stiffness of the basket that we and others (31, 48) measured and is also supported by a recent study which describes the molecular source of this flexibility in yeast (71). The other scenario is a rapid exchange of entire baskets or individual filaments. Tpr-depleted NPCs are rarely observed in normal conditions - assessed by fluorescent labeling from us (Figure 4A) and others (25, 72). However, individual filaments may rapidly exchange from NPCs and reassemble in different configurations. This is supported by data showing the high exchange rate of Tpr and Nup153 (21–23). In particular, Nup153 dynamics may destabilize Tpr association to the scaffold and loosen filament attachment. This is in agreement with many of our individual NPC images where the basket appears asymmetrical both in terms of topography and Tpr localization (Figure 3). It was also shown recently in budding yeast that peripheral NPC components are loosely linked to the main scaffold and their interactions with the pore are labile (22). In this study the authors suggest that the dynamic nucleoporins could be envisioned as “a cloud of accessory factors surrounding and constantly exchanging with the NPC in vivo”. Such a model could explain how the basket filaments could rapidly change conformation while exchanging. This rapid switching could be a source of high variability in basket conformation and account at least in part for the heterogeneity we measure.

Although our findings challenge the textbook image of an ever protruding basket, the fact that this structure can explore the central channel is functionally relevant. Indeed, the basket serves as a docking platform for mRNAs sorting and export (73–76). The capacity of the basket filaments to explore around the pore and their extended shape makes great assets to scan the pore vicinity for mRNA. As a matter of fact Tpr depletion reduces the docking frequency of mRNAs to the nuclear pore and affects their diffusion through the central channel (76, 77). Interestingly, the last steps of mRNA export and mRNP remodeling occur at the cytoplasmic filaments (78). These filaments were often thought of as extended structures outreaching to the cytoplasm orthogonally from the NPC. However, it has been shown in yeast that the Nup82 complex, a major constituent of these filaments, in fact stands over the central channel (79). This surprisingly showed that the filaments also have the ability to bend towards the central channel. But it also explains how mRNPs are continuously handled from the central channel, where they interact with FG-repeat Nups through transport receptors, to their final maturation site at the cytoplasmic filaments. Our findings about the flexibility of the NPC basket and its ability to explore the central channel thus complete this picture and propose a continuous path for mRNPs during their journey across the NPC.

In a more general context, the permeability barrier paradox has fascinated biologists and physicists for decades: how can a single structure allow active transport of extremely large particles while retaining non-eligible molecules above 5 nm in size? This ability necessarily has to do with a high degree of plasticity in the interaction surfaces between pore components and cargoes. Our present work is a first step towards understanding the dynamics and adaptation capacity of the NPC; this will certainly be extended in the coming years with developments in correlative high-speed AFM.

## Materials and Methods

### Cell strains, antibodies and siRNA

U2OS cells were obtained from ATCC. CRISPR-engineered Nup96-SNAP U2OS cell lines (*47*) were obtained from CLS (clsgmbh.de). siRNAs were ordered from Eurogentec: siNup153 according to the sequence in (*80*) (AAGGCAGACUCUACCAAAUGUUU dTdT) and scrambled siRNA (UAGAUACCAUGCACAAAUCC dTdT).

All antibodies and fluorescent labels used in this study are commercially available. The purchase references and dilutions used are listed in Table 1.

**Table 1:**
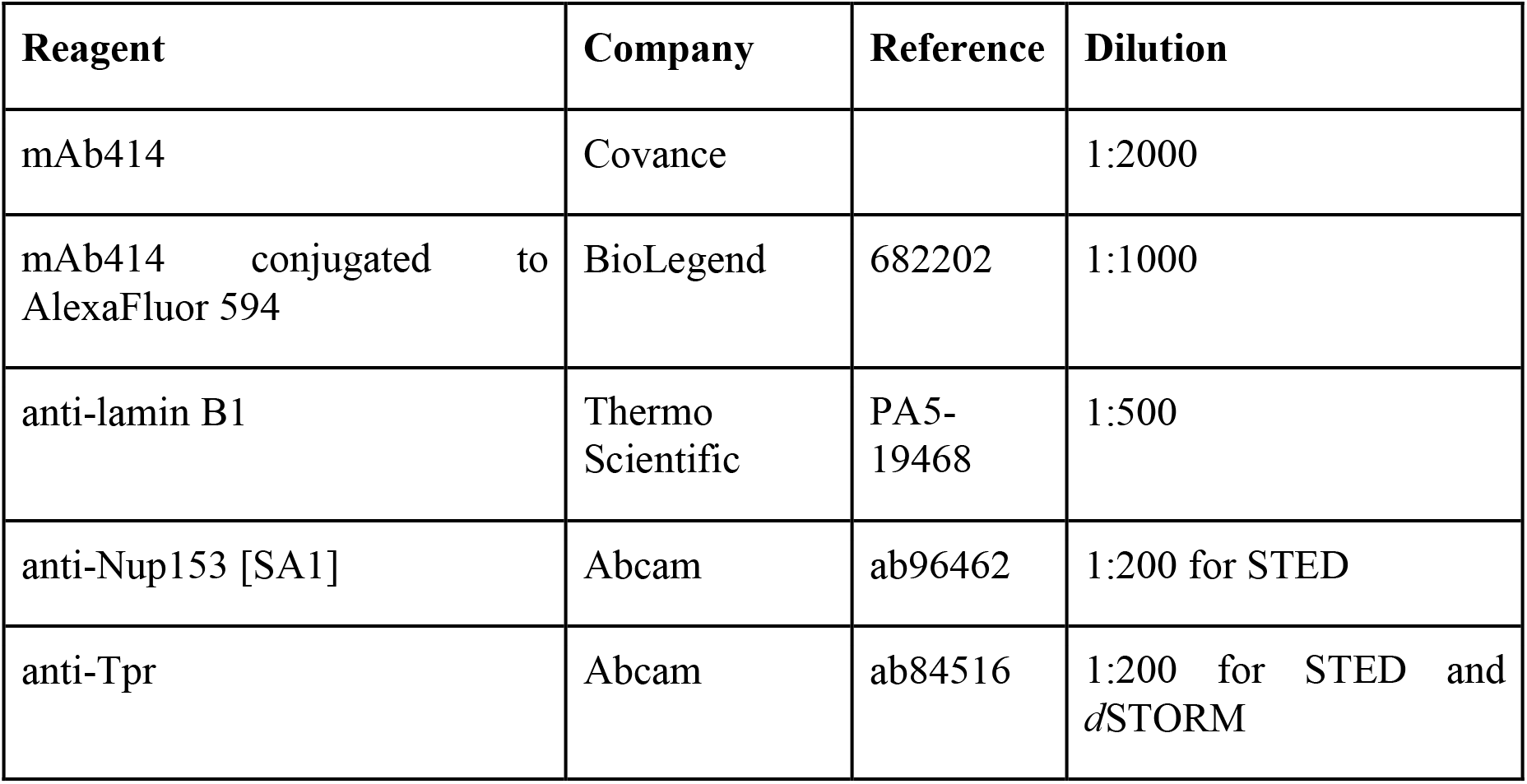
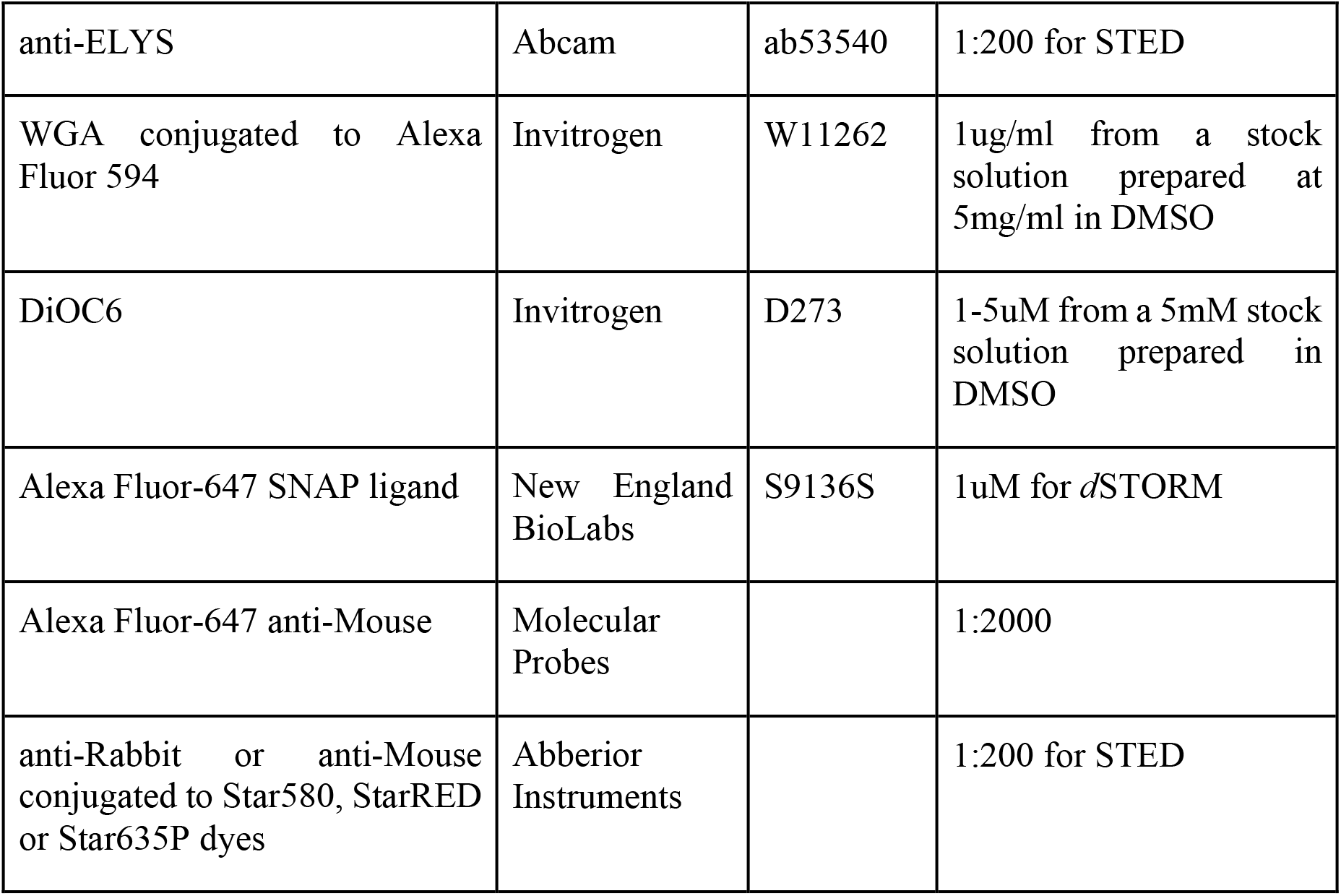
Reagents for fluorescent labeling.

### Cell culture and transfection

Cells were cultured in DMEM / Glutamax (Gibco) supplemented with 10% Fetal Calf Serum (Gibco) in a humid atmosphere at 37°C, 5% CO2.

For siRNA transfection, 1.5 millions of cells were seeded in 10cm diameter plates. The day after they were transfected with 1.25 nmol of siRNA and 10uL of Lipofectamine 2000 (Invitrogen) according to the manufacturer’s recommendation. Medium was changed after 4-6 hours. Transfection was repeated 48h later in identical conditions. Cells were lysed or imaged the day after.

### Preparation of nuclear envelopes for AFM

All buffer compositions are indicated in table 2

**Table 2:** Buffers.

| Buffer | Composition | Reference |
| --- | --- | --- |
| Hypotonic buffer | 50mM Tris pH7.5, 1mM DTT, Complete (Roche) | Ori *et al*, 2013 (46) |
| S500 | 10mM HEPES pH7.5, 2.5mM MgCl<sub>2</sub>, 50mM KCl, 500mM Sucrose, 1 mM DTT, Complete | Doucet *et al*, 2010 (57) |
| NE-A | 0.1mM MgCl<sub>2</sub>, 1mM DTT + 30U DNaseI (Sigma, D5319) + 250U RNase A | Ori *et al*, 2013 (46) |
| NE-B | 20mM Tris pH8.5, 10% sucrose, 0.1mM MgCl<sub>2</sub>, 1mM DTT | Ori *et al*, 2013 (46) |
| Nuclei buffer | 10mM HEPES pH7.5, 2.5mM MgCl<sub>2</sub>, 83mM KCl, 17mM NaCl |  |
| Immunofluorescence (IF) buffer | PBS wit 10mg/ml BSA, 0.1% Triton X-100, 0.02% SDS | Doucet *et al*, 2010 (57) |
| Permeabilization buffer (PB) | PBS with 0.4% Triton X-100 | Thevathasan *et al*, 2019 (47) |
| Quenching solution (QS) | 100mM NH<sub>4</sub>Cl in PBS | Thevathasan *et al*, 2019 (47) |
| dSTORM buffer | PBS supplemented with 10% glucose, 0.5 mg/mL glucose oxidase (Sigma), 0.04 mg/mL catalase (Sigma, C3556), 35mM mercaptoethylamine (MEA). | Thevathasan *et al*, 2019 (47) |

Nuclei were prepared from cultured cells as previously described (*46*). Briefly, around 40 millions cells (four 75cm^2^ confluent flasks) were trypsinized. Cells were rinsed in PBS then hypotonic buffer (HB) by 5 minutes centrifugation at 300g. Cells were then resuspended in 1mL HB, incubated on ice for 45 minutes and disrupted by mechanical shearing using a Dounce with a tight pestle (approximately 10 strokes are sufficient). Cell lysis was controlled by visual inspection using an inverted microscope (Primovert, ZEISS). The nuclei suspension was then loaded on top of 5mL of S500 and spun for 15 minutes at 300g. The supernatant containing cellular debris was discarded, the pellet was resuspended in 300ul of S500, transferred to a clean tube containing 5mL of S500 and spun for 3 min at 300g. The pellet, containing mostly intact cells, was discarded. The supernatant was transferred to a clean tube and spun for 15 minutes at 300g. Nuclei were resuspended in S500 at a final density of 10^6^ nuclei per mL.

25mm high-precision type 1.5 glass coverslips of 0.17mm thickness (Marienfeld) were coated with 250uL of Poly-L-Lysine (Sigma) diluted 1:10 in water and filtered (0.01mg/ml final concentration) for 2 minutes at room temperature. 250uL nuclei were deposited on dried coverslips and incubated for 15 minutes at room temperature, the suspension was then removed and the coverslip was rinsed with S500.

Meanwhile, the Nuclear Envelope buffers NE-A and NE-B (Table 2) were pre-warmed at 30°C. Nuclei were treated with 200ul of NE-A for 30 seconds. 800ul of NE-B were added and the whole mixture was incubated for 30-40 minutes at room temperature as described in (*46*). The nuclease mixture was removed and nuclei were rinsed several times with S500.

### Labeling for confocal, STED and correlative TIRF/AFM microscopy

Cells and nuclei were labeled as previously described (*57*). Briefly, they were rinsed in PBS and fixed with 4% PFA (Electron Microscopy Sciences) for 5 minutes, then rinsed twice in PBS. For intact cells labeling, an additional permeabilization step was performed for 10 minutes in IF buffer (Table 2). To keep the nuclear envelope intact, nuclei were simply incubated in blocking buffer (PBS supplemented with 3% goat serum) for 10 minutes. Primary antibodies were diluted in blocking buffer and incubated for 1 hour. Samples were rinsed 3 times in PBS. Secondary antibodies were incubated for 30 minutes; samples were then rinsed 3 times in PBS. All incubations were performed in the dark at room temperature. For confocal microscopy, coverslips were mounted on glass slides on a drop of Vectashield and sealed with nail polish. Slides were kept at 4°C until imaging. For AFM microscopy, samples were kept at 4°C in S500 until mounting in a coverslip holder designed to be adapted to JPK AFM and based on AttoFluor (Thermofisher) geometry.

### Labeling for *J*STORM

25mm high-precision type 1.5 glass coverslips of 0.17mm thickness (Marienfeld) were sequentially washed in acetone, ethanol and water, then sonicated in 1M KOH for 20 minutes. Coverslips were then extensively washed in milliQ water and air dried. For intact cells, the coverslips were UV-sterilized before seeding. 24h later, cells were washed 2x with PBS and prefixed in 2.4% PFA for 30sec. Cells were then permeabilized with PB (Table 2) for 3 minutes, and fixation was completed in 2.4% PFA for 30 additional minutes. The fixing agent was quenched in QS buffer (Table 2) for 5 minutes, rinsed twice with PBS for 5 min and blocked for 10 min with PBS + 0.1% BSA.

Staining with SNAP-ligand was performed for 2h. Samples were then washed 3 times with PBS for 5 min, and post-fixed in PBS + 4% PFA for 5 minutes. All incubations were performed at room temperature. For imaging, the coverslips were set in an AttoFluor cell chamber and covered with 1.2 mL of dSTORM buffer (Table 2), covered by a glass coverslip to limit oxidation. The buffer was renewed after 2 hours.

### Atomic Force Microscopy

Atomic force microscopy was performed on a JPK NanoWizard 4 combined with a CellHesion Z stage (Bruker Nano GmbH, Germany) and a homemade *d*STORM setup (*28*). Experiments were performed in liquid in Nuclei buffer, derived from Kramer et al. (*68*). Bruker MSNL probes using cantilever E or Olympus OTR4 probes (with respective nominal stiffnesses 0.1 and 0.12 N/m) were used for open nuclei. Both MSNL and OTR4 probes have pyramidal tips with a nominal tip radius of 2 nm for MSNL and 7 nm for OTR4. The spring constant of the cantilever was calibrated, prior to imaging, by acquisition of a force versus distance curve on a clean glass coverslip. Optical lever sensitivity was calibrated with a linear fit of the repulsive part of the force curve. Then, the spring constant was finally evaluated using the thermal noise method (*81*).

Images were recorded at resolution of 256 x 256 pixels in Quantitative Imaging (QI) mode. The force setpoint was below 300 pN to minimize the force applied while keeping the acquisition time short enough to avoid drift and facilitate the correlation with the superresolution image. The length of the force curve was 200-300 nm to avoid tip-sample adhesion and facilitate tip detaching in each indentation cycle, with a tip approach-retract speed of typically 30 μm/s.

### Fluorescence microscopy

Confocal images were acquired on a Leica SP8 microscope with a 63x/1.4 NA Plan Apo objective. STED images were acquired on an Abberior Instrument Expert Line STED super-resolution microscope (Abberior Instruments GmbH, Göttingen, Germany), using 561- and 640-nm pulsed excitation laser sources and a pulsed STED laser operating at 775 nm and an 80-MHz repetition rate. The fluorescence excitation and collection were performed using a 100x/1.40 NA Plan Super Apochromat objective (Olympus, Hamburg). All acquisition operations were controlled by Imspector software (Abberior Instruments GmbH, Germany). Image series were recorded using Imspector software with the following parameters: pixel dwell time, 3.3 μs; 10 repetitions; pixel size, 15 nm, STED power (measured in the back focal plane of the objective), 250-300mW. The STED laser was pulsed with a gap delay of 700 ps and a duration of 8 ns.

*d*STORM experiments were performed on an AFM/*d*STORM correlative setup (*28*) consisting of a JPK NanoWizard 4 coupled with an inverted microscope equipped with an oil immersion Zeiss α-Plan-Apochromat 100x, 1.46 NA DIC objective. Proteins of interest in cells and nuclei were labeled with AlexaFluor 647.

An oxygen-scavenging *d*STORM buffer containing 100 nm size Tetraspeck (Molecular Probes #T7279, 1/1000 dilution) fluorescent beads was loaded on the sample. A volume of 1.2 mL of this solution was used to fill the AttoFluor chamber. A glass coverslip was put on top of the AttoFluor to limit oxygen exchange with the ambient atmosphere.

*d*STORM acquisition was performed by illuminating the sample with a 642 nm laser, set to 500 mW nominal power to obtain 3-7 kW/cm^2^ on the sample. 30 000 frames, with an acquisition time per frame of 20-30 ms, were recorded with an Andor iXon Ultra 897 emCCD 512×512 pixels camera, with a pixel dimension of 16 μm. A 2x telescope was placed in front of the camera to obtain a final image magnification of 200. 0-0.1 kW/cm2 of 405 nm was sometimes used for conversion from the dark state and to facilitate the dye blinking. The acquisition area was chosen so that a nucleus surrounded by several Tetraspeck fiducial markers was visible.

### Data analysis

Images were treated with Fiji.

*d*STORM analysis was performed using the Fiji plug-in Thunderstorm(*82*). Drift was corrected thanks to the fiducial markers or in some cases by cross-correlation proposed by the plugin. To reconstruct the image, localizations were filtered to keep only intensity (number of photons) between 500 and 5000, localization precision below 45 nm and standard deviation of the fit of the Point Spread Function (PSF) between 90 and 180 nm. For diameter measurements, analysis was performed using SMAP (Ries, 2020).

AFM images were treated using Gwyddion software. Images were corrected by subtracting the mean plane, applying line correction and sometimes removing horizontal lines. Aberrant pixels (very high or very low) due to problems in the force curve were sometimes corrected by using a mask which interpolates a pixel with surrounding pixels. For crops around NPCs, data from images with scan sizes of 1 to 3 μm were used and crops of 300 nm x 300 nm were extracted. The Z scale was always ranging from 0 to 150 nm which is sufficient for the topography around NPCs. The offset and tilt of the nucleoplasmic ring was corrected using the Matlab algorithm from Stanley et al. (*31*). To calculate the rotationally averaged height profiles, we measured the height profile along a 150 nm line crossing the pore center. We then rotated the line of 0.05 π Rad and repeated the process 20 times (figure S2A), scanning the entire structure; the resulting radially averaged profiles are then plotted and fitted with three gaussians corresponding to the scaffold rim and the basket distal ring (Figure S2C). The distance between the two extreme gaussian peaks corresponds to pore diameter. Pore depth is calculated as the vertical distance between the average ring height and height at the pore center. This was automatically performed thanks to custom scripts written in Python.

For mechanical measurements from QI data, we used JPK data processing software. Experimentally calibrated spring constant was used for calculations, force curve baseline offset and tilt was subtracted before tip vertical position correction. Tip-sample contact point was determined using the Hertz/Sneddon model for a quadratic pyramid with tip pyramid angles modified to match MSNL probe geometry. The stiffness was measured by fitting the repulsive part of the force from the contact point over 20 nm. We did not generate Young modulus maps since its calculation is influenced by the tip-sample contact geometry that could vary when imaging very corrugated samples.

## Acknowledgments

We thank Jean-Bernard Fiche, Antoine Le Gall and Marcelo Nollmann for their help and advice regarding SMLM. We thank the members of the team ‘Integrative Biophysics of Membranes’ for their continuous support and feedback (https://integrativebiophysicsofmembranes.wordpress.com). The science-art model in figure 6 was designed and drawn by Zhanna Santybayeva (illustration4science.com).

## Author contributions

AV performed all AFM and AFM-correlative experiments; LC was involved in analysis of AFM force curves; PD was involved in AFM data analysis; PR and GB were involved in sample preparation and optimization; HH helped design and implement AFM experiments on nuclear envelope samples; OF helped design, optimize and analyze dSTORM data; CD performed confocal and STED experiments, wrote scripts for data analysis; PEM and CD designed and supervised the project. The manuscript was written by CD, AV and PEM and proofread by all authors.

## Data and materials availability

All data, code, and materials used in the analyses will be available to any researcher for purposes of reproducing or extending the analyses.

