## Supplemental material for "Structure and mechanics of the human Nuclear Pore Complex basket"

**This PDF file includes:**

Supplementary Text
Figs. S1 to S5

**Discussion about the mechanical assessment of NPCs**

The stiffness values given by Stanley *et al*. are higher than what we measured in human NPC, both at the location of the nucleoplasmic ring and basket. Several parameters can explain this difference, including the tip size and geometry, and also the fact that PeakForce QNM used by Stanley *et al*. probe the NPC at much higher loading rate than in our study and thus estimate an elastic response at higher frequency (typically kHz for PeakForce), naturally stiffer, compared to our quasi-static frequency range (typically 10-100 Hz for QI). Indeed, supplementary control experiments performed by Stanley *et al*. using Force Volume, which probe the sample at lower speed, exhibit much lower stiffness values, closer to our results.

It thus appears that the basket, which is already very soft, is suspended on top of an even softer medium. Thus, we wondered if the basket could be deflected by the AFM tip during imaging which would affect the topographical images. Indeed, height was measured in most cases at a force of 240 pN, which is low for AFM imaging, but high compared to NPC stiffness. We thus reconstructed the sample height with different forces applied (Figure S5B): height images reconstructed at 60, 120, 180 and 240 pN show no significant differences in their overall topographies. We can therefore exclude that the force applied leads to an inversion in topography. Nevertheless we see a small indentation, and thus the height we measure might be slightly underestimated. However, the distribution of conformations that we describe remains valid and truly revisits the textbook view of NPC basket organization.



-Bruker Confidential-

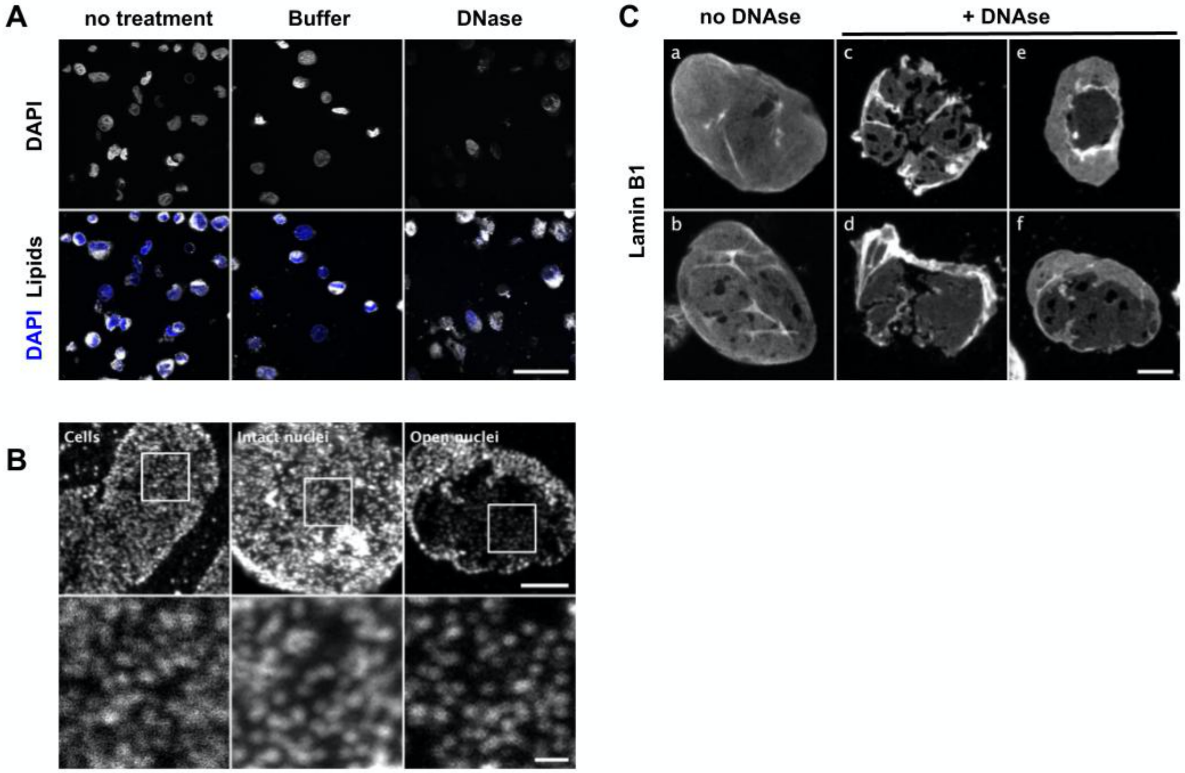


**Fig. S1. Supplementary information for figure 1**

A- Confocal images of nuclei extracted from U2OS and treated or not with nucleases or buffer. Samples were labeled with Hoechst (blue) and a lipid dye (DiOC6, grey scale) and imaged by confocal microscopy. Scale bar is 50 µm. B- Confocal images of cells, nuclei and nuclear envelopes labeled with mAb414. The boxed areas are zoomed in the lower panel. Scale bars are 5 µm (top) and 1 µm (bottom). C- Confocal imaging of nuclei treated (c-f) or not (a-b) with nucleases and labelled with anti-lamin B1. Well-circumscribed openings (e) better preserve membrane integrity, as depicted by homogenous lamin staining; scale bar is 5 µm.



-Bruker Confidential-

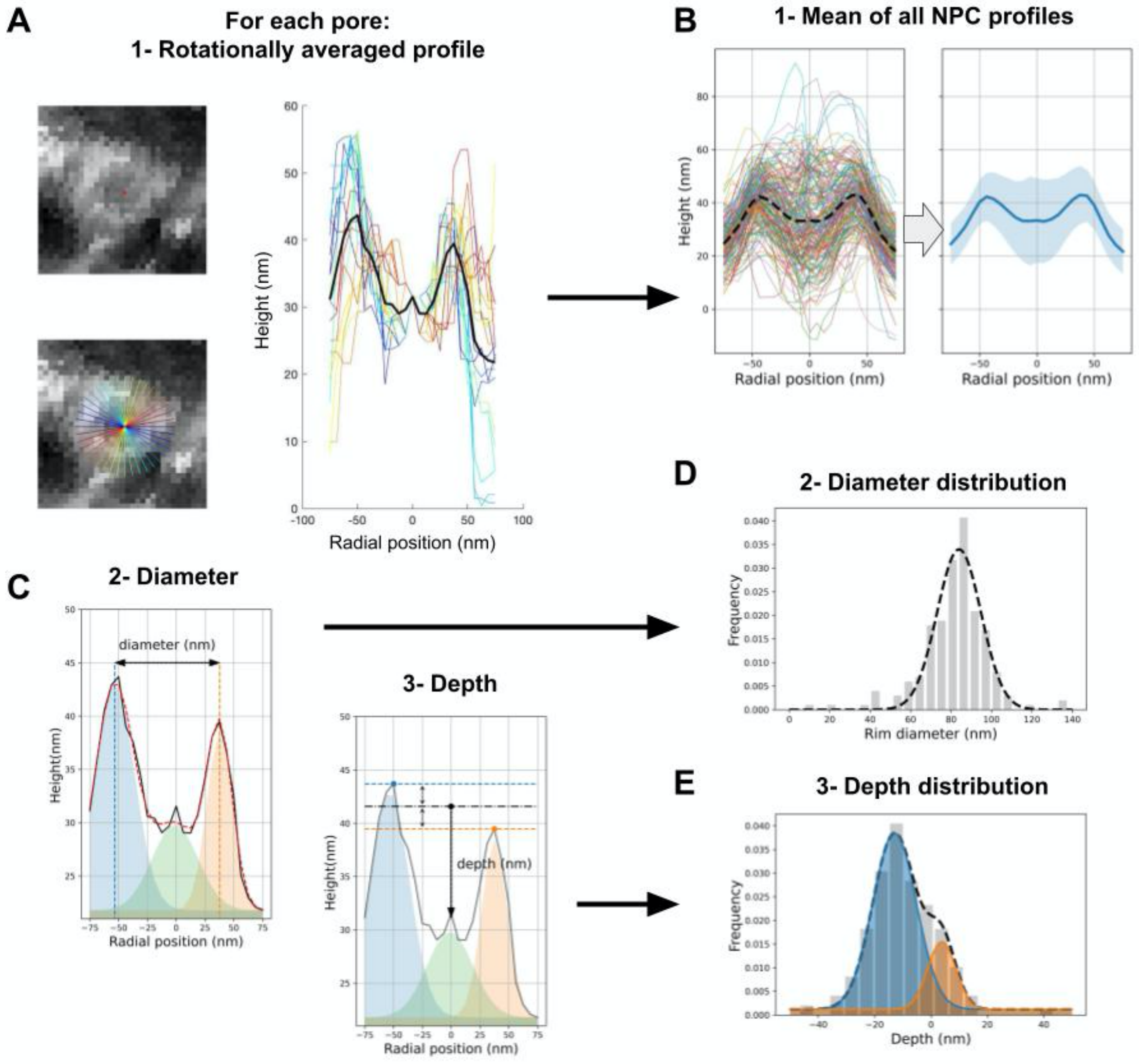


**Fig. S2. Supplementary information for figure 2**

A- Individual pores are manually cropped from 2µm x 2µm AFM scans. For each pore, a height profile is measured along a 150 nm-line drawn across pore center. The line is then rotated 0.05 π up to π and a profile is generated at each angle. The 20 profiles are then averaged to produce a single rotationally averaged height profile per pore (black line). B- The profiles of >200 pores were averaged to show the mean height profile of NPCs (right panel, the shaded area corresponds to the standard deviation). C- Pore diameter and depth are calculated from individual rotationally averaged height profiles: each profile is fitted with three gaussians; the diameter is the distance between the first and last gaussian peaks (corresponding to the ring position). The depth is calculated as the height difference between the pore center (radial position = 0) and the average ring height. Depths are thus positive in protruding pores and negative in collapsed ones. D-E- Diameter and depths distributions of >200 NPCs are then plotted as frequencies and fitted with one (respectively two) gaussians. The dashed line represents the fit function.



-Bruker Confidential-

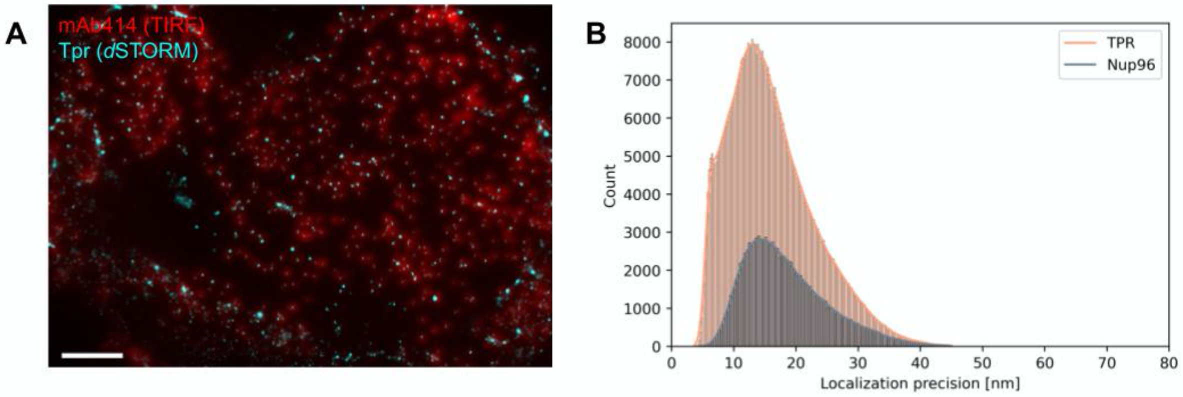


**Fig. S3. Supplementary information for figure 3**

A- Composite image of a NE labeled with mAb414 coupled to Alexa Fluor 594 and anti-Tpr detected with an anti-Rabbit coupled to Alexa Fluor 647.The TIRF image of the 594 channel is merged with the reconstructed map of the *d*STORM acquisition in the 647 channel. Scale bar is 2µm.

B - Distribution of STORM localization precision for TPR and Nup96 labelled with Alexa Fluor 647. Data were extracted from 2 typical images.



-Bruker Confidential-

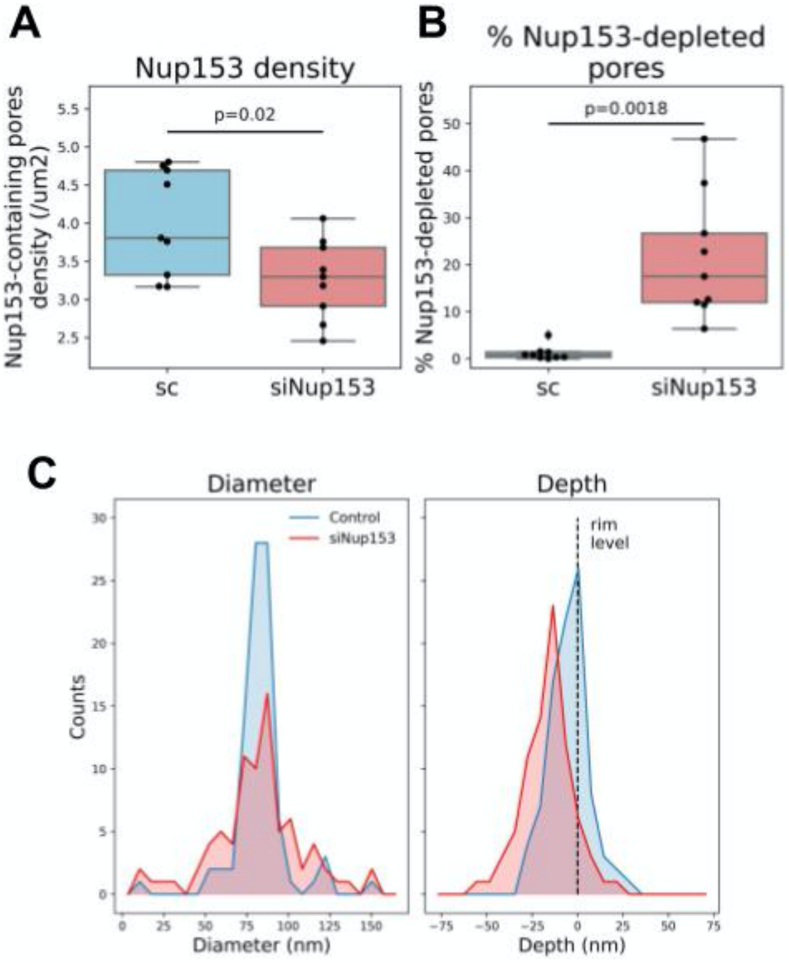


**Fig. S4. Supplementary information for figure 4**
U2OS/GFP-Nup96 cells were treated with scrambled or Nup153-specific siRNAs. The density of pores positively labeled with anti-Nup153 was counted for 9 nuclei in each condition (A). From the same nuclei we calculated the percentage of pores (= Nup96+) with no detectable Nup153 labeling (B).
C- NPC diameter and depth were measured from rotationally averaged height profiles of over 80 pores from control or Nup153-depleted nuclei.



-Bruker Confidential-

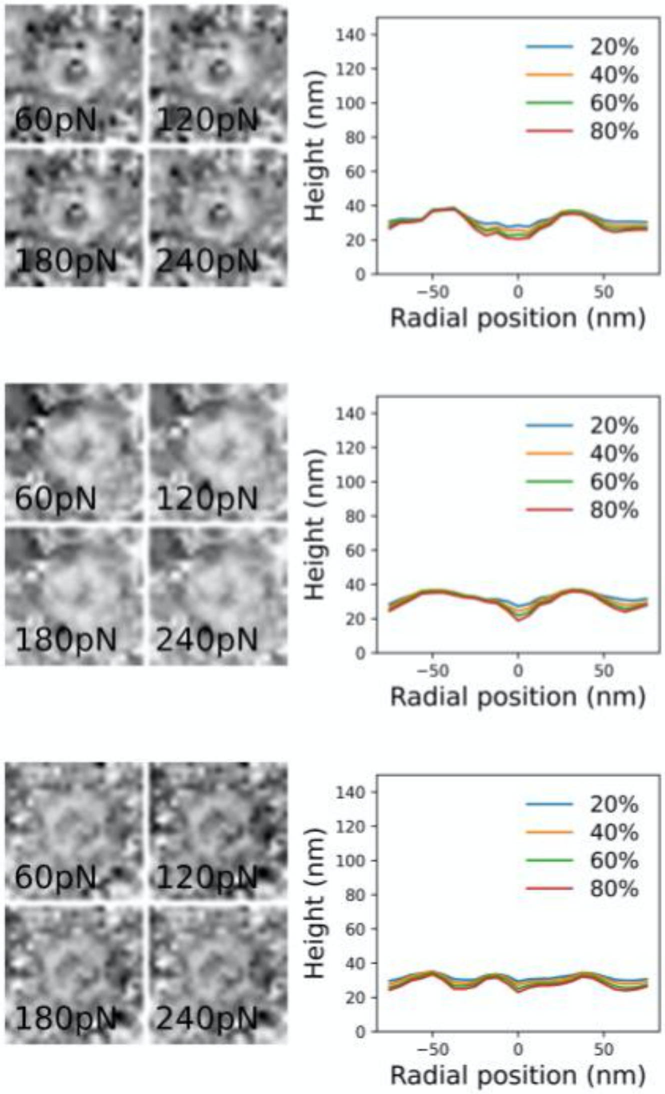


**Fig. S5. Supplementary information for Figure 5**
Topography images of individual pores reconstructed at 60, 120, 180 and 240 pN with their corresponding rotationally averaged height profiles.



-Bruker Confidential-
